# Drinking microstructure analysis of direct controls of water intake in male and female rats

**DOI:** 10.64898/2025.12.30.697026

**Authors:** Bo W. Sortman, Emalie L. Mullane, Kennedy L. Lamb, Andrea A. Edwards, Ann-Marie Torregrossa, Derek Daniels, Jessica Santollo

## Abstract

Measures of rodent drinking (lick) patterns are used to test how various manipulations affect fluid intake. Decades of research indicate that manipulating the satiating potency of a stimulus affects the number of licking bursts whereas manipulating the hedonic value of the solution affects the size of the licking bursts or rate of licking. This type of analysis has been largely applied to manipulations that modulate drinking that is already occurring (i.e., indirect controls or modulators of intake), but there is a dearth of studies with detailed measures of licking in response to stimuli that are primary drivers of intake (i.e., direct controls). A better understanding of licking behavior in response to direct controls of fluid intake could provide insight into how drinking is stimulated. Accordingly, we analyzed lick patterns after varying doses of primary drivers (direct controls) of intake, angiotensin II, hypertonic saline, or carbochol, in male and female rats. As expected, all treatments dose-dependently increased licks for water. Analysis of licking patterns revealed dose-related increases in burst number after treatment with all stimuli, but dose did not predict burst size or lick rate. Together with previous work on licking patterns these data suggest that direct controls of water intake are more related to processes associated with satiety than to changes in hedonic value of the consumed substance.

## INTRODUCTION

Drinking (lick) microstructure analysis is a powerful tool used to analyze patterns of fluid (water, saline) and liquid food (sucrose, ensure, etc.) intake. This approach can provide insight about the mechanisms underlying any observed changes in ingestion. Although there are slight methodological differences between lab groups, researchers examining short term water or saline intake tests typically analyze patterns of bursts, which are often defined as at least 2 licks with an interlick interval less than 1 second [1–3]. Foundational work by Smith and Davis demonstrated that changes in the number of drinking bursts within a test session reflect changes in negative feedback from post-oral sites, whereas the average size of those bursts (licks/burst) and the rate of licking reflect changes in orosensory or hedonic value of the substance being consumed [4–6]. Applying this to water intake, however, may require modification in terminology because accumulating evidence demonstrates the importance of the oral cavity in thirst termination [7–10]. Nevertheless, negative or terminating feedback, whether or not it is gastric in origin, plays a role in the number of bursts. Therefore, measuring licking patterns during drinking can shed light on mechanisms underlying ingestion.

Recently, we developed a framework, using terminology that was inspired by the classical work from Smith [11], to categorize different bioregulators (hormones, neurotransmitters, neuromodulators) as controlling fluid intake directly or indirectly [12]. According to this framework, a direct control is a bioregulator that causes the animal to switch from a state of not drinking to a state of drinking. Treatment with the hormone Angiotensin II (AngII), which potently and rapidly stimulates drinking is an example of a direct control [13–16]. In turn, an indirect control is a bioregulator that affects the likelihood that drinking continues or terminates. The hormone estradiol is an example of an indirect control. It modulates the drinking that is already happening, but does not cause a rat to initiate drinking on its own [17–24]. Although decades of research have used drinking microstructure analysis to probe for changes in water and saline intake after manipulations to indirect controls, the literature lacks a systematic evaluation of the differences in lick patterns that underlie changes in total intake when varying the dose of the direct stimulus for intake. The absence of this analysis made it unclear if changes in intake related to manipulations of direct controls of intake would be a function of burst size, burst number, or both.

The scarcity of detailed measures of licking in response to treatments that act as direct controls of drinking is perhaps because microstructure analysis was not a common practice during the early studies that meticulously characterized different drinking stimuli. Furthermore, recent studies rarely, if ever, vary the strength of the direct control. As such, if lick patterns were analyzed, these tests would find changes in all aspects of microstructure because the comparison is to an animal that is not drinking at all. To answer this long-standing unexplored question in the literature, we performed experiments to test for systematic changes in drinking microstructure in dose-response studies of different direct controls of water intake. These experiments serve multiple purposes. The primary purpose is to evaluate the licking patterns that accompany direct controls of intake. For example, dose-related increases in burst size or lick rate would suggest that a direct control increases intake, at least in part, because it increases the positive feedback associated with the consumed water. Alternatively, if burst number is the only measure that changes with increasing doses of a direct control, it would suggest that drinking initiation and maintenance are driven by the lack of satiating feedback. Additionally, these data will allow us to explore differences in the licking patterns induced by multiple stimuli that our framework considers to be in the same category. In this respect, consistency in the observed pattern would add support to the framework by demonstrating alignment of things the framework deems alike.

## METHODS

### Animals

Male and female Sprague Dawley rats (Envigo/Inotiv Laboratories, Indianapolis, IN) were used in all studies described. For Experiments 1 and 3, rats were singly housed in modified hanging wire-mesh stainless steel cages with *ad libitum* access to food (Teklad 2018; Harlan Laboratories) and tap water unless otherwise noted. For Experiment 2, rats were singly housed in modified shoebox cages with *ad libitum* access to food (Teklad 2018; Harlan Laboratories) and tap water unless otherwise noted. All testing occurred in the rats’ home cages during the early part of the light phase. The temperature- and humidity-controlled colony rooms were maintained on a 12:12 h light-dark cycle (lights on at 0700 h). All experimental protocols were approved by the Animal Care and Use Committee at the University of Kentucky or the University at Buffalo, and the handling and care of the animals was in accordance with the *National Institute of Health Guide for the Care and Use of Laboratory Animals*.

### Surgery

Animals in Experiments 1 and 3 underwent stereotaxic surgery to implant a chronic cannula aimed at the right lateral ventricle following standard laboratory procedures. Briefly, rats were anesthetized with an intramusclar injection of a ketamine [i.m., 75 mg/kg (males), 80 mg/kg (females); Fort Dodge Animal Health, Fort Dodge, IA] and xylazine [i.m., 5 mg/kg (males), 4.5 mg/kg (females); Akron Inc., Decatur, Il]. The rat’s head was shaved and secured into a stereotaxic frame. A small incision was made on top of the skull, a small hole was drilled, and a 26-gauge guide cannula was implanted using the following coordinates: 0.9mm posterior and 1.4mm lateral to bregma, and 2.8mm ventral to the skull. The cannula was fixed to the skull with bone screws and dental cement. All rats received a single injection of carprofen (sc, 5 mg/kg; Pfizer Animal Health, New York, NY) after surgery to minimize pain and a single injection of isotonic saline (sc, 5 ml) to replace any lost fluids. Five to seven days later, accurate cannula placement was verified by measuring the drinking response to an injection of 10 ng Ang II. Only rats that drank at least 5 ml in 20 min after Ang II-treatment were included in the study.

### Intake Measures

Total intake was calculated by weighing the water bottles before and after each test period. Licking behavior was measured using a contact lickometer (designed and constructed by the Psychology Electronics Shop, University of Pennsylvania, Philadelphia, PA for Experiments 1 & 3/University of Kentucky, College of Arts and Sciences Electronics Shop for Experiment 2) that recorded individual licks to allow for analysis of drinking microstructure. The lickometer interfaced with a computer using an integrated USB digital I/O device (National Instruments, Austin, TX). Home cages were affixed with an electrically isolated metal plate with a 3.175-mm-wide opening, through which the rat needed to lick to reach the drinking spout, minimizing the possibility of non-tongue contact with the spout. A drinking (lick) burst was defined as at least 2 licks with an inter-lick-interval (ILI) of no more than 1 s and burst size was defined as the number of licks within a burst, as previously described [1, 2]. The rate of licking was determined using the parameters established by [25]. The rate of licking for the test period was calculated by dividing the total number of licks in all bouts by the duration of all bouts, with a bout being defined as at least 2 licks with an inter-lick-interval of no more than 5 minutes. Lick rates were then normalized to licks/60 seconds.

### Experiment 1

Lick microstructure analysis for AngII dose-response. To address this, we performed a *de novo* analysis from a previously published data set from our lab [26]. Using a counterbalanced crossover design, age-matched male (n = 7) and female (n = 7) rats received intracerebral ventral (i.c.v.) injections of 0, 1, 10, 50, or 100ng AngII dissolved in 1 µl tris buffered saline (TBS). Water intake during the subsequent 2 h was monitored. Food was removed from the cages during the 2 h test period. All females were tested during the estrus stage of the estrous cycle. Testing occurred once every four to five days until all rats were tested with every dose of AngII.

### Experiment 2

Lick microstructure analysis for hypertonic saline dose response. Using a counterbalanced crossover design, age-matched male (n = 6) and female (n = 6) rats received a 0.1 ml sc injection of lidocaine and then were immediately injected with 1 ml of 0.15. 0.5, 1 or 2 M NaCl (sc). Rats were returned to their home cages, food and water were removed, and pre-weighed water bottles were provided 30 min later. Water intake during the subsequent 2 h was monitored. Testing occurred once a week until all rats were tested with every dose of hypertonic saline.

### Experiment 3

Lick microstructure analysis for carbachol dose response. Age-matched male (n = 10) and female (n = 10) rats received intracerebral ventral (i.c.v.) injections of 0, 0.1, 1, 10, or 100ng carbachol dissolved in 1 µl tris buffered saline (TBS). Water intake during the subsequent 2 h was monitored. Food was removed from the cages during the 2 h test period. Testing occurred once every three days. Due to headcap/cannula failure, not all rats received all drug doses, therefore a different statistical analysis was used for this experiment (see Data Analysis).

### Data Analysis

All data are presented as means ± SEM throughout. All experiments used a two-way mixed design with sex (male/female) as the between-subjects factor and dose (AngII, hypertonic saline, carbachol) as the within-subjects factor. A two-way mixed ANOVA was used except in the case of missing values, where a mixed-effects model was fit using Restricted Maximum Likelihood (REML) (burst size analysis in Experiment 1 and 2, and all analysis in Experiment 3). Newman-Keuls post hoc tests were used to follow up main or interactive effects.

## RESULTS

### Experiment 1

Lick microstructure analysis for AngII dose-response. AngII dosage had a significant effect on intake, licks, and drinking microstructure. As shown in Table 1, volume (ml) consumed was influenced by a main effect of sex [F(1,12) = 7.57, p < 0.05] a main effect of dose [F(4,48) = 128.99, p < 0.001] and an interaction between dose and sex [F(4,48) = 5.71, p < 0.001]. Males consumed more water than females (p < 0.05). AngII dose-dependently increased water intake, but intake after treatment with 50 and 100ng were not significantly different from each other (p < 0.05). Males consumed more water than females after treatment with 10 and 100ng AngII (p < 0.01). A similar pattern was found for total licks during the test period. As shown in Figure 1, licks for water were influenced by a main effect of sex [F(1,12) = 7.55, p < 0.05], a main effect of dose [F(4,48) =63.00, p < 0.001], but the interaction between dose and sex was not statistically significant [F(4,48) = 2.47, p = 0.057]. Males had more licks for water than females (p < 0.05). Licks for water after 0 and 1 ng AngII were less than 10-100ng AngII. Licks after 10ng AngII were less than licks after 100ng AngII. Licks after 50ng AngII were greater than 0-1ng AngII but not different from 10 and 100ng AngII. Licks after 100ng AngII were greater than licks after 0-10ng AngII (p < 0.05). Burst number was influenced by a main effect of dose, [F(4,48) = 30.76, p < 0.001], but was not influenced by sex [F(1, 12) = 2.42, p = n.s.], and the test for an interaction between sex and dose was likewise not significant [F(4,48) = 1.17, p = n.s.; Figure 1B]. Burst number after treatment with 10-100ng AngII was greater compared to burst number after treatment with 0 or 1ng AngII (p < 0.05). Burst number after treatment with 50 or 100ng AngII was greater than burst number after treatment with 10ng AngII (p < 0.05). There was no effect of sex [F(1,12) = 0.1692, p = n.s.] or dose [F(2.286, 26.86) = 2.026, p = n.s.], and the interaction between sex and dose was not significant [F(2.286, 26.86) = 0.03, p = n.s.], on burst size (Figure 1C). The rate of licking was influenced by a main effect of dose [F(2.273, 26.71) = 6.89, p < 0.01], but there was no main effect of sex [F(1,12) = 0.72, p = n.s.] or an interaction between sex and dose [F(2.273, 26.71) = 0.94, p = n.s.; Figure 1D]. The rate of licking after treatment with 10, 50, and 100ng AngII was significantly less than the rate of licking after vehicle treatment (p < 0.05).

**Figure 1.**
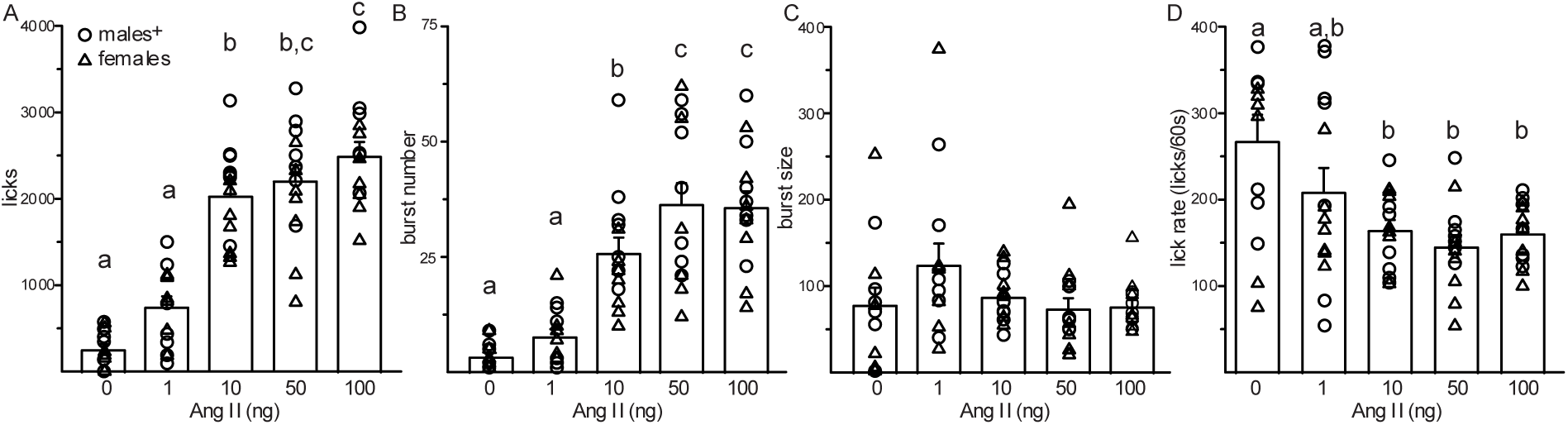
AngII dose dependently increased water intake in male and female rats. (A) Doses dependent increases in licks were observed. Licks for water were greater after 10, 50, and 100ng AngII compared to 0 and 1ng AngII. Licks for water after 100ng AngII was greater than licks after 10ng AngII. Males had more licks for water compared to females. (B) Dose dependent increases in burst number were observed. Burst number was greater after 10ng AngII, compared to 0 and 1ng AngII. Burst number after 50 and 100ng AngII was greater than burst number after 0, 1, and 10ng AngII. (C) There were no differences in burst size as a function of AngII dose. (D) The rate of licking after 10, 50, and 100ng AngII was less than the rate of licking compared to 0ng AngII. +Males > females, p < 0.05. Different letters denote significant treatment differences, p < 0.05.

**Table 1.**
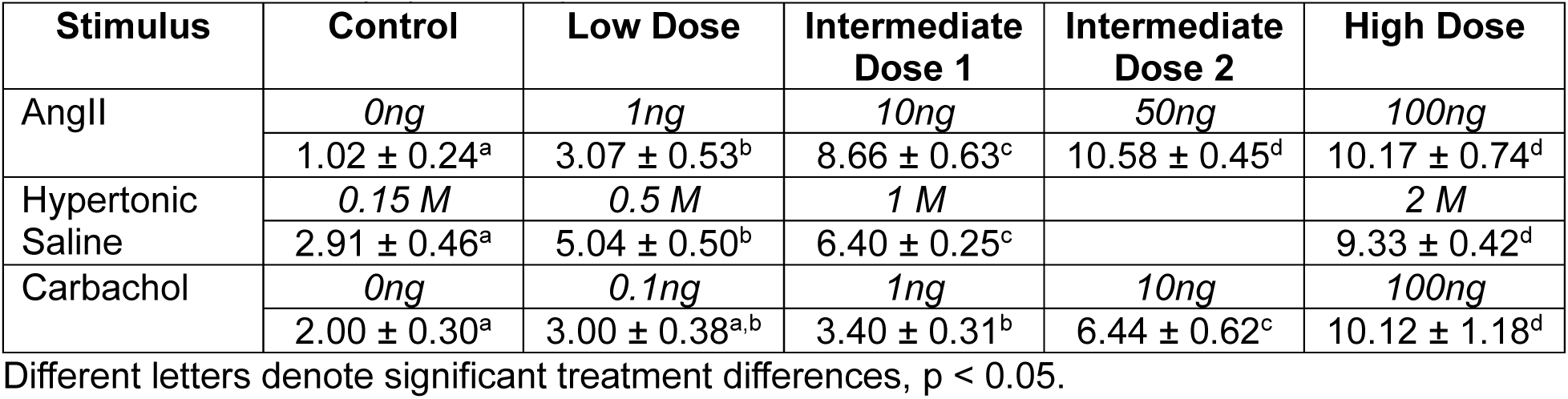
Water intake (ml) consumption.

### Experiment 2: Lick microstructure analysis for hypertonic saline dose response

Hypertonic saline dosage had a significant effect on intake, licks, and drinking microstructure. Volume (ml) consumed was influenced by a main effect of dose [F(3,30) = 81.08, p < 0.001], but there was not a main effect of sex [F(1,10) = 1.91, p = n.s.], or an interaction between sex and dose [F(3,30) = 0.87, p = n.s.; Table 1]. As the dose of hypertonic saline increased, water intake significantly increased between each dose. An identical effect was observed with total licks. Licks for water were influenced by a main effect of dose [F(3,30) = 44.90, p < 0.001], but there was not a main effect of sex [F(1,10) = 2.02, p = n.s.] or an interaction between sex and dose [F(3,30) = 0.90, p = n.s.; Figure 2A]. As the dose of hypertonic saline increased, water intake significantly increased between each dose. Burst number was influenced by a main effect of dose [F(3,30) = 6.35, p < 0.005], but there was not a main effect of sex [F(1,10) = 0.35, p = n.s.] or an interaction between sex and dose [F(3,30) = 1.81, p = n.s.; Figure 2B]. Burst number after treatment with 2M NaCl was greater than treatment after 0.15 and 0.5 M NaCl. Burst number after treatment with 1M NaCl was not significantly different from any treatment. Burst size was influenced by a main effect of dose [F(2.039, 19.71) = 5.85, p < 0.005], but there was not a main effect of sex [F(1,10) = 0.6436, p = n.s.] or an interaction between sex and dose [F(2.039, 19.71) = 2.332, p = n.s.; Figure 2C]. Burst size after 0.5, 1, and 2 M NaCl was greater than after control (0.15M) treatment. The rate of licking was influenced by a main effect of dose [F(2.507, 24.24) = 6.62, p < 0.01], but not by a main effect of sex [F(1,10) = 2.99, p = n.s.] or by an interaction between sex and dose [F(2.507, 24.24) = 0.24, p = n.s.; Figure 2D]. The rate of licking after treatment with 0.5, 1, and 2M NaCl was significantly greater than the rate of licking after vehicle treatment (p < 0.05).

**Figure 2.**
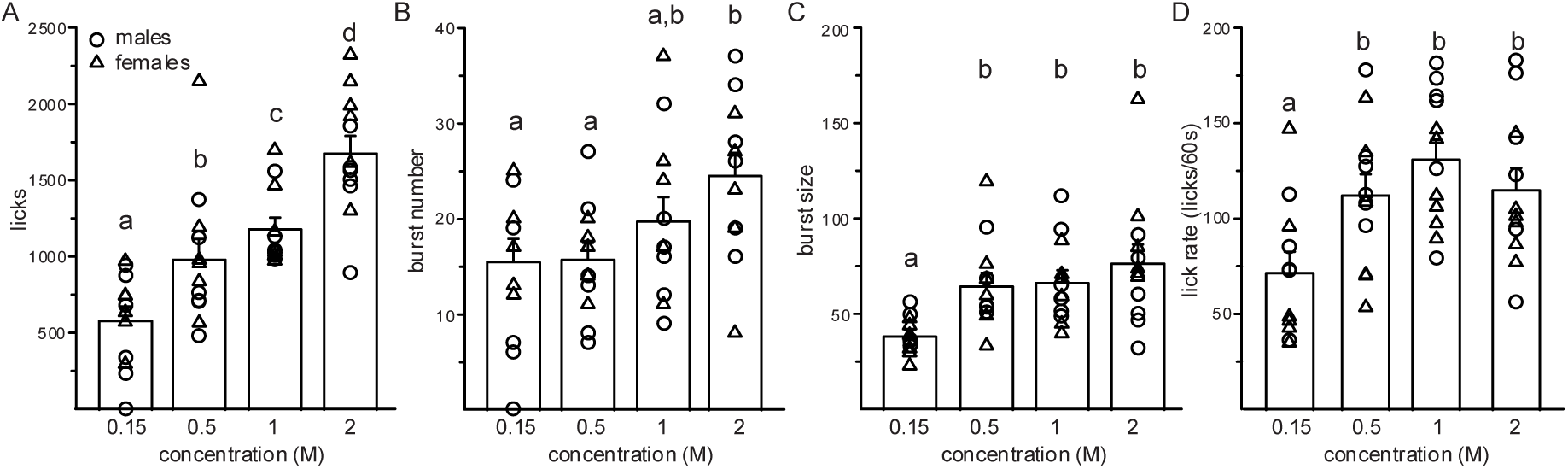
Hypertonic saline dose dependently increased water intake in male and female rats. (A) Dose dependent increases in licks were also observed after treatment with 0.5, 1, and 2 M NaCl. (B) Treatment with 2 M NaCl increased burst number compared to treatment with 0.5 and 1 M NaCl. (C) All doses of hypertonic saline increased burst size. (D) All doses of hypertonic saline increased lick rate compared to control treatment. Different letters denote significant treatment differences, p < 0.05.

### Experiment 3: Lick microstructure analysis for carbachol dose response

Carbachol dosage had a significant effect on intake, licks, and lick microstructure. Volume (ml) consumed was influenced by a main effect of sex [F(1,18) = 6.53, p < 0.05] and dose [F(2.15,24.73) = 30.23, p < 0.0001], and by an interaction between dose and sex [F(4,46) = 5.33, p < 0.005; Table 1]. Males consumed more water than females (p < 0.05). While water intake was not different between 0 and 0.1ng carbachol, treatment with 1ng carbachol was greater than control but not different than intake after 0.1ng carbachol. Intake after 10ng carbachol was greater than 0-1ng treatment and intake after 100ng carbachol was greater than all other doses. Males consumed more water than females after treatment with 100ng AngII (p < 0.005). A similar pattern was found for total licks during the test period. Licks for water was influenced by a main effect of sex, [F(1,18) = 6.59, p < 0.05], by a main effect of dose [F(2.25,25.83) = 32.26, p < 0.0001], and by an interaction between dose and sex [F(4,46) = 4.03, p = 0.01; Figure 3A]. Males had more licks for water than females (p < 0.05). Licks for water after 0.1 and 1ng carbachol were greater than control. Licks for water after 10ng carbachol were greater than 0-1ng treatment and licks after 100ng carbachol were greater than all other doses. Burst number was influenced by a main effect of dose [F(1.34,15.45) = 11.35, p < 0.005], but there was not a main effect of sex [F(1,18) = 0.08, p = n.s.] or a significant interaction between sex and dose [F(4,46) = 0.55, p = n.s.; Figure 3B]. Burst number after 0.1 and 1ng carbachol were greater than control. Burst number after 10ng carbachol was greater than 0-1ng treatment and burst number after 100ng carbachol were greater than all other doses. There were not main effects of sex [F(1,18) = 1.02, p = n.s.] or dose [F(1.53,17.22) = 2.75, p = n.s.], or an interaction between sex and dose [F(4,45) = 1.19, p = n.s.], on burst size (Figure 3C). The rate of licking was not influenced by a main effect of dose [F(2.596, 28.56) = 0.88, p = n.s.], or by a main effect of sex [F(1,18) = 1.98, p = n.s.], and the interaction between sex and dose was also not significant [F(2.596, 28.56) = 1.57, p = n.s.; Figure 3D].

**Figure 3.**
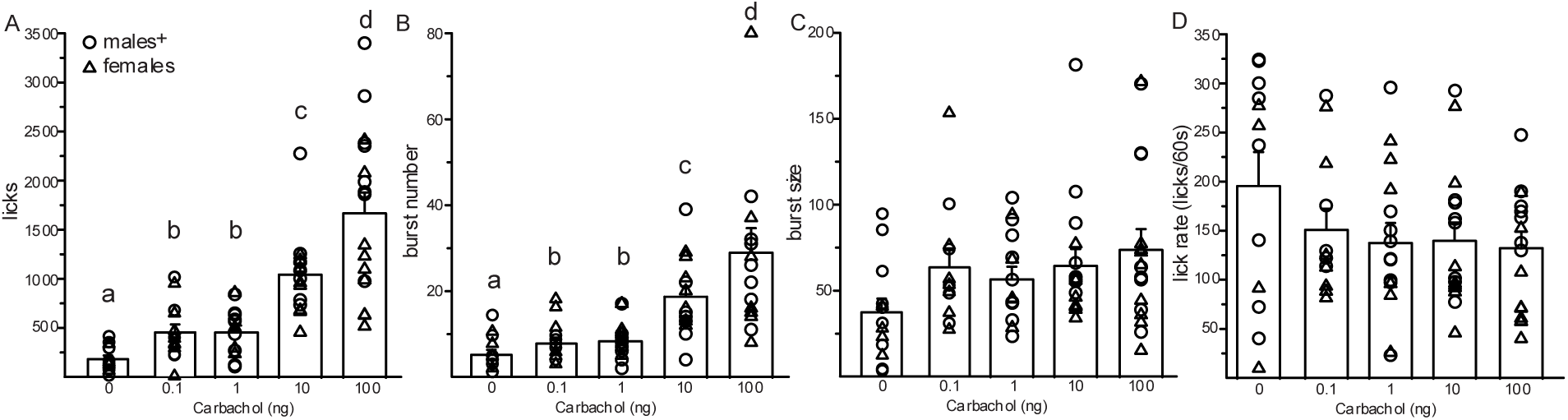
Carbachol dose dependently increased water intake in male and female rats. (A) Dose related increases in licks for water were observed. Licks after 0.1 and 1ng were greater than control but not different from each other. Licks for water after 10ng carbachol were greater than 0-1ng and licks for water after 100ng carbachol were greater than 0-10ng. Males had more licks for water compared to females. (B) Dose related increases in burst number were observed. Burst number after 0.1 and 1ng were greater than control but not different from each other. Burst number after 10ng carbachol were greater than 0-1ng and burst number after 100ng carbachol were greater than 0-10ng. (C) There were no differences in burst size as a function of carbachol dose. (D) There were no differences in the rate of licking as a function of carbachol dose. +Males > females, p < 0.05. Different letters denote significant treatment differences, p < 0.05.

## DISCUSSION

Analysis of drinking microstructure patterns have yielded helpful insight into the mechanisms underlying changes in fluid intake and provide direction of future research questions. For example, drinking microstructure analysis of the fluid inhibitory effects of estradiol demonstrate that the decreased intake is the result of changes in burst number, implicating changes in satiety signals as a mechanism [17, 27, 28]. This led us to investigate interactions between estradiol and glucagon-like peptide-1 (GLP-1), a drinking satiety signal. Indeed, we found that estradiol enhanced the fluid inhibitory effect of the GLP-1 agonist, exendin-4, suggesting that an interaction between these two hormones may underlie the anti-dipsogenic effect of estradiol [17]. Surprisingly, a review of the literature found minimal, if any, drinking microstructure analysis for direct controls of fluid intake, aka a stimulus that causes an animal to initiate drinking. This analysis may have been previously dismissed as not useful because an animal going from not drinking to drinking would always have an increase in all microstructure measures as the comparison would be to zero (no bursts, no burst size). If, however, there are systematic changes in drinking microstructure after introduction of a direct control of intake, it could provide mechanistic insight into how drinking is initiated. Accordingly, we asked if there are uniform changes in drinking microstructure patterns in response to direct controls of water intake as a function of the dose of the stimulus. In support of our hypothesis, we found dose-related increases in burst number, but not in burst size or in rate of licking, following treatment with each of the three stimuli tested: AngII, hypertonic saline, and carbachol. These results fill a gap in the literature and suggest that the drinking that occurs after initiation by a direct control is a function of either overcoming or modulating the strength of satiety, and that continued drinking after initiation by a direct control is a function of the satiating potency of the consumed substance.

AngII is the key hormone that stimulates drinking during hypovolemia. A drop in either blood pressure or osmolality triggers the synthesis of AngII which binds to its receptor, angiotensin type 1 receptor, in the lamina terminals to stimulate water (and saline) intake. As expected, we observed dose-related increases in licks for water. A similar dose-related pattern was observed with burst number. Burst size, however, was not different between doses and there was no difference in the rate of licking between any of the doses of AngII that significantly increased licks. Others have reported that increased doses of Ang decrease the latency to drink [29] which could be interpreted as an increase in motivation to initiate fluid intake and intraoral infusion of AngII, which isolates the consummatory phase of ingestion, increases latency to drip compared to vehicle [30]. To the best of our knowledge, the present report is the first to provide a detailed analysis that is focused on the effect of AngII on licking microstructure in rats. Thus, this conspicuous absence has been corrected by the present studies, and the data indicates that water intake simulated by AngII drives bursts of licking, rather than the length or rate of licking bursts.

Next, we examined drinking microstructure patterns as a function of dose of hypertonic saline treatment. Hypertonic saline treatment induces osmotic dehydration, due to hyperosmolality of the extracellular fluid. This assault to fluid homeostasis induces water intake, with saline/salt being actively avoided. We observed dose-dependent increases in licks for water after treatment with hypertonic saline. We also observed dose-related increases in burst number, with the highest dose of hypertonic saline (2M) increasing burst number over treatment with 0.5M and the 1M dose having an intermediate effect. Burst size, however, did not differ by dose, nor did the overall rate of licking. To our knowledge there are no published reports examining drinking microstructure, including latency to drink, following treatment with varying doses of hypertonic saline. These results mimic our findings from Expt. 1 and suggest that these direct controls act by driving licking bursts, rather than by increasing the time between the initiation and termination of a burst.

Finally, we examined drinking microstructure patterns as a function of carbachol dose. Carbachol is a non-selective cholinergic agonist that increases thirst dependent on muscarinic Gi/o activity [31]. Injections of carbachol directly into the subfornical organ or median preoptic nucleus stimulate drinking [32, 33]. Additionally, ICV injections of carbachol increase Fos expression in the Organum Vasculosum of the Laminae Terminalis [34]. As expected, we observed dose-related increases in licks for water after treatment with carbachol. The dose-related differences in licks was a function of burst number, but not related to burst size or rate of licking. Again, to our knowledge there are no published reports examining any aspect of drinking microstructure after treatment with carbachol. Similar to our findings in Expt. 1 and 2, these data suggest that the strength of the stimulus to drink is associated with the number of bursts, but not the size of the bursts.

AngII, hypertonic saline, and carbachol all exhibited dose related changes in burst number, but not burst size or rate of licking. The consistency in the effects amongst what we had considered to all be direct controls of intake supports the proposed framework, but does this allow us to draw conclusions about the mechanism by which drinking is initiated? There was no change in burst size or rate of licking as a function of dose of AngII, hypertonic saline, or carbachol. This suggests that the increased drive to drink is not associated with an increase in the hedonic value of the water. In other words, the drinking that occurs once initiated does not appear to be sustained by the palatability of the water, and the water does not appear to be a source of positive feedback. Instead, our data suggest that direct controls of intake are associated with decreased satiating potency of the fluid consumed. This could suggest that the initiation of drinking involves overcoming a state of satiety. While satiety conceptually is a lack of thirst, mechanistically this may be different than a lack of hunger. Indeed, previous work on eating behavior suggests that active neural and hormonal signals are required to maintain satiety [35–38]. In contrast, no such signals have been found to maintain a lack of thirst. Instead, thirst arises from an insult to fluid homeostasis that requires correction [39, 40]. These data point to a hypothesis that can now be tested. As the drive to drink increases, are stronger satiety signals needed to affect fluid intake? Follow up studies will be needed to test this idea, but this example highlights the benefit of this type of analysis: it provides clues about the mechanisms of the controls of intake, providing direction for future research.

Sex differences were observed in the drinking response to AngII and carbachol, but not in the response to hypertonic saline. Males consumed more water and had more licks for water after treatment with AngII and carbachol. We reported this sex difference in AngII-stimulated water intake in our previous report which used this dataset [26, 41], which corroborates earlier findings demonstrating that males consume more water in response to AngII than females [42]. This sex difference is mediated, at least in part, by the fluid inhibitory effects of estradiol [19, 20, 23, 42, 43]. To our knowledge, this is the first report of sex differences in carbachol’s dipsogenic effect, however, sex differences in other physiological responses to carbachol have been reported. For example, carbachol has a greater effect in male rats, compared to females, at stimulating vasopressin release [44]. This sex difference in the dipsogenic effect of carbachol, however, may not be mediated by circulating estrogens, as a previous report found that estradiol failed to reduce carbachol-stimulated drinking in ovariectomized female rats [42]. Here we found that males and females consumed similar amounts of water after treatment with hypertonic saline. This confirms a previous report from our lab [45] and supports previous research demonstrating that estradiol does not influence osmotic thirst [20, 21, 46, 47].

In conclusion, this series of experiments demonstrates that increased strength of a drinking stimulus is associated with changes in burst number, but not burst size or overall lick rate. This suggests that the drinking that is initiated by a direct control is a function of changes in satiety signals independent of the physiological nature of the dipsogen (osmotic vs hypovolemic). These nuances revealed by the analyses can be tested by future research, demonstrating the power of this approach to provide clues into the mechanisms underlying changes in intake and generating further experimental questions. Indeed, 30 plus years after its introduction to the literature, drinking microstructure analysis remains a useful tool for the refined analysis of ingestive behaviors.

## Acknowledgements

This work was supported by the National Science Foundation [grant 2019346] to JS and the National Institutes of Health [grant DK133818] to DD and National Science Foundation [grant 1942291] to A-MT.

## Notes

### Competing Interest Statement

The authors have declared no competing interest.

